# CMiNet: An R Package and User-Friendly Shiny App for Constructing Consensus Microbiome Networks

**DOI:** 10.1101/2025.05.09.653200

**Authors:** Rosa Aghdam, Claudia Solís-Lemus

**Affiliations:** Wisconsin Institute for Discovery, University of Wisconsin-Madison, Madison, WI; Department of Plant Pathology, University of Wisconsin-Madison, Madison, WI

**Keywords:** Compositional data, Microbial interactions, Microbiome networks, Consensus network construction, Biomarker discovery

## Abstract

1. Microbial networks offer critical insights into community structure, ecological interactions, and host–microbe dynamics. However, constructing reliable microbiome networks remains challenging due to variability among existing inference methods, limited overlap between networks, and the absence of a gold standard for validation.
2. We developed `CMiNet` (https://cminet.wid.wisc.edu), an interactive Shiny application and R package that enables consensus microbiome network construction by integrating up to ten widely used inference algorithms. `CMiNet` supports both correlation-based and conditional dependence-based methods and provides users with flexible options to construct individual or consensus networks across different approaches.
3. `CMiNet` employs a consensus-based filtering strategy that retains only edges supported by multiple methods, leading to more robust network structures that better reflect underlying biological interactions. This approach enhances reproducibility, minimizes method-specific biases, and improves biological interpretability. Its user-friendly interface allows researchers to upload data, customize network construction parameters, visualize networks interactively, and export results—without requiring programming expertise.
4. We demonstrated `CMiNet` using both gut and soil microbiome datasets. In the soil microbiome application, robust network construction enabled the consistent identification of disease-associated taxa. Specifically, we identified a set of key taxa, including *Ktedonobacteria, Acidobacteriae, Vicinamibacteria, MB-A2-108, Planctomycetes*, and *Anaerolineae*, selected by multiple independent methods.

## 1. INTRODUCTION

Microbial communities play vital roles in host health, ecosystem functioning, and environmental resilience [1, 2]. To understand the structure and dynamics of these communities, researchers often turn to microbiome co-occurrence networks, where nodes represent microbial taxa and edges represent inferred interactions or associations. Recent advances in network inference methods, including equation-free approaches, have enabled researchers to reconstruct complex ecological interaction networks from observational microbiome data [2]. These networks are increasingly used to uncover ecological relationships, identify potential keystone species, and guide therapeutic interventions [3, 4, 5]. Developing robust computational frameworks for analyzing complex biological communities, including microbial networks and broader biodiversity datasets, remains an essential frontier in ecological research [6]. In this context, constructing accurate microbial networks from high-throughput sequencing data is essential for uncovering patterns of co-variation among taxa and for revealing ecological or functional dependencies within microbial communities. These inferred networks are widely applied in studies of gut microbiota and disease, soil health, host–microbe interactions, and environmental microbiomes [7, 8, 9]. However, inferring reliable networks remains challenging due to data-specific complexities such as compositionality, sparsity, and high dimensionality, all of which can introduce spurious associations and reduce reproducibility across studies [10, 11]. To address these methodological challenges, researchers have developed a wide range of network inference algorithms, including correlation-based methods such as SparCC [12]; as well as conditional dependence-based approaches including `SpiecEasi` [13], SPRING [14], and our recently developed CMIMN algorithm [15], which leverages conditional mutual information to capture both linear and non-linear dependencies. Although these methods can identify relationships in complex microbiome networks, selecting a single algorithm often leads to networks that may vary significantly, raising concerns about the reliability of the inferred interactions [5]. Such variability arises from the different assumptions and statistical frameworks each method employs, often resulting in limited overlap among inferred networks. This discrepancy motivates the development of more integrative approaches. The lack of a gold standard for validating microbiome networks further complicates their interpretation. Unlike gene regulatory networks, which can occasionally be benchmarked against established biological pathways [16, 17], microbial interactions are unknown, variable across environments, or highly context-specific. This uncertainty hinders the ability to make robust biological conclusions and limits cross-study comparability [18]. These limitations have fueled growing interest in generating more reliable and interpretable networks using consensus approaches. Consensus network construction—by integrating outputs from multiple algorithms—can help reduce method-specific bias and improve biological relevance. However, consensus construction is often a labor-intensive process, requiring users to manually run several tools, align outputs, and harmonize edges across networks. This makes it inaccessible for many researchers, especially those without programming experience.

To bridge this gap, we first developed `CMiNet`, an R package that integrates ten widely used microbial network inference algorithms into a unified framework. The package allows users to construct consensus networks by integrating results from either correlation-based methods, conditional independence-based methods, or a combination of both. To make this framework accessible to a broader research community, we then created a user-friendly Shiny application that eliminates the need for programming or package installation. Both the R package and the Shiny app feature intuitive interfaces for selecting algorithms, tuning parameters, constructing consensus networks, and visualizing network structures across varying confidence thresholds. We describe the design and implementation of `CMiNet`, demonstrate its application using benchmark datasets, and provide guidance for its use in ecological and biomedical research. By enhancing accessibility, reproducibility, and robustness, `CMiNet` supports broader adoption of network-based microbiome analysis across disciplines. One powerful application of constructing microbiome networks is the comparison of microbial communities across different conditions, such as healthy versus diseased states. This type of analysis can reveal how network structure, connectivity, and taxon centrality shift in response to environmental or physiological changes. However, drawing meaningful conclusions from such comparisons requires robust and reproducible network inference. `CMiNet` addresses this need by enabling the construction of more reliable microbiome networks. First, we apply `CMiNet` to construct separate networks for healthy and diseased samples and then compare them using topological features and taxon-specific centrality metrics, such as Influence-Vertex Importance (IVI). This comparative analysis allows us to identify differentially central taxa, uncover potential microbial biomarkers, and better understand how microbial communities respond to disease.

## 2 METHODS

### 2.1 Overview of consensus network construction

The core of the `CMiNet` framework is its ability to construct a consensus microbiome network by integrating results from multiple inference algorithms. The input to `CMiNet` is a microbiome abundance matrix, where rows represent samples and columns correspond to microbial taxa. Users can select from a variety of algorithms and specify parameters for each. Each selected algorithm infers a microbial network independently. These individual networks are then merged into a weighted network, where the edge weight reflects the number of algorithms that support a given interaction between taxa. A final consensus network is generated by applying a user-defined threshold; only edges supported by a specified number of algorithms are retained. This approach increases confidence in inferred microbial relationships by focusing on reproducible patterns. Figure 1 provides a detailed overview of the `CMiNet` workflow for consensus microbiome network construction. The process begins with a microbiome abundance matrix, where rows represent samples and columns represent microbial taxa. This matrix serves as the input for all selected network inference algorithms. Users choose one or more network inference methods and set algorithm-specific parameters. Each selected method infers a microbial interaction network independently. `CMiNet` combines the results from selected inference algorithms to construct a weighted consensus network, where the edge weight reflects the number of algorithms that support a given interaction between microbial taxa. For example, the edge between Microbe 1 and Microbe 2 is supported by all ten methods and is therefore assigned a weight of 10. In contrast, the edge between Microbe 4 and Microbe 5 is supported by only two methods, resulting in a weight of 2. To refine the network and focus on robust interactions, users specify a threshold that filters out low-confidence edges. Only interactions supported by at least the chosen number of algorithms are retained in the final network. For example, if the threshold is set to 8, the edge between Microbe 1 and Microbe 2 (weight = 10) is retained, while the edge between Microbe 4 and Microbe 5 (weight = 2) is excluded.

**Figure 1.**
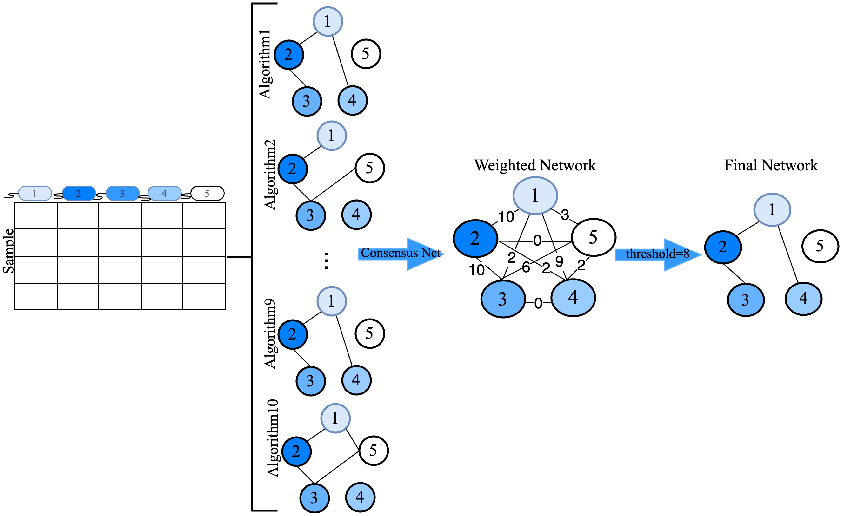
Overview of the CMiNet workflow. Users input a microbiome abundance matrix, select inference algorithms, and construct a weighted network where edge weights indicate the number of supporting methods. A threshold is applied to retain high-confidence edges. For instance, an edge with weight 10 (supported by all methods) is retained when the threshold is set to 8, whereas an edge with weight 5 is excluded.

### 2.2 User-defined inference modes in CMiNet

To accommodate the diversity of the microbiome research goals and the methodological differences among inference algorithms, `CMiNet` offers three user-defined modes for network construction. This flexibility allows researchers to build networks based on either correlation-based methods, conditional independence-based methods, or a combination of both, enabling tailored analyses aligned with the study context. The *correlation-based methods* option includes only algorithms that estimate marginal associations between microbial taxa. These methods, such as Pearson [3], Spearman [4], Bicor [4], and SparCC [12], identify co-occurrence patterns without controlling for indirect interactions or confounding variables. While computationally efficient, these approaches may capture indirect associations and are sensitive to compositional effects. The *conditional independence-based methods* mode uses algorithms that estimate direct interactions by accounting for the influence of other taxa in the network. Methods such as `SpiecEasi` (MB and Glasso) [13], SPRING [14], GCoDA [19], and CMIMN [15] aim to recover more biologically meaningful relationships by inferring conditional dependencies, often resulting more interpretable networks. The *consensus networks* mode combines both algorithm types to construct a consensus network. In this mode, edges are retained only if they are supported across multiple selected methods—regardless of type—thus providing higher confidence in the inferred microbial interactions. This integrated approach reduces method-specific bias and supports more reproducible network analysis.

### 2.3 CMiNet R package and Shiny app

The `CMiNet` R Package provides a programmatic environment for constructing consensus microbiome networks. Users can access all ten inference algorithms, adjust their parameters, and generate consensus networks directly within the R environment. The package includes utility functions for visualization, thresholding, and exporting networks in standard formats for downstream analysis. All codes and documentation are available on Github https://github.com/solislemuslab/CMiNet. The `CMiNet` Shiny App ensures accessibility for users without programming experience by combining algorithmic power with a user-friendly design. It lowers technical barriers and broadens the usability of consensus microbial network inference across diverse research backgrounds. All code and documentation are available on GitHub at https://github.com/solislemuslab/CMiNetShinyAPP. The `CMiNet` Shiny app is freely available at https://cminet.wid.wisc.edu.

### 2.4 Overview of network inference algorithms and their parameters

To support flexible and robust analysis, `CMiNet` includes ten widely used microbial network inference algorithms. Each algorithm has distinct modeling assumptions, strengths, and limitations, making it important for users to tailor their choice of methods and parameters to the characteristics of their microbiome dataset. To facilitate this, `CMiNet` allows full control over method-specific parameters in both the R package and the Shiny App. Table 1 summarizes the inference models included in `CMiNet`, outlining their category, core modeling approach, key advantages, known limitations, and the default parameters used in our implementation.

**Table 1.** Summary of microbial network inference methods implemented in CMiNet.

| Method | Model/Approach | Advantages | Limitations | Default Parameters |
| --- | --- | --- | --- | --- |
| Pearson [20] | Linear correlation | Simple and fast; widely used | Sensitive to outliers; assumes normality and linearity | None |
| Spearman [21] | Rank-based correlation | Robust to outliers; captures monotonic relationships | Ignores compositional bias; limited to pairwise relations | None |
| Bicor [4] | Biweight midcorrelation | Down-weights outliers; suitable for noisy data | Limited adoption; similar limitations to Pearson/Spearman | None |
| SparCC [12] | Correlation of log-ratios assuming sparsity | Handles compositional data; robust to violations of sparsity; provides interpretable correlations via log-ratio transformation | Computationally intensive for large datasets; may yield non-positive definite covariance matrices | imax = 20, kmax = 10, alpha = 0.1, Vmin = 1e-4 |
| SE_MB [13] | Neighborhood selection via MB algorithm (graphical lasso) | Infers conditional dependencies; robust to compositionality; works in high-dimensional settings | Sensitive to regularization parameter; limited performance with excess zeros | method = 'mb', lambda.min.ratio = 1e-2, nlambda = 15, rep.num = 20, ncores = 4 |
| SE_glasso [13] | Sparse inverse covariance estimation (graphical lasso) | Produces sparse, interpretable networks; robust to compositionality; suitable for high-dimensional data | Computationally intensive; assumes Gaussianity; may not capture non-normal features of microbiome data | method = 'glasso', lambda.min.ratio = 1e-2, nlambda = 15, rep.num = 50 |
| SPRING [14] | Semi-parametric rank-based partial correlation | Handles zero-inflated data; robust to misspecification of total abundance; estimates direct interactions | May miss non-linear relationships; does not account for compositionality; computationally intensive | Rmethod = 'original', quantitative = TRUE, nlambda = 15, rep.num = 20, ncores = 5 |
| gCoda [19] | Penalized likelihood under logistic normal model (GLM) | Adjusts for compositional bias; supports differential analysis of networks | Sensitive to pseudo-count choice; assumes GLM structure; non-convex optimization may lead to local minima | counts = FALSE, pseudo = 0.5, lambda.min.ratio = 1e-4, nlambda = 15, ebic.gamma = 0.5 |
| CCLasso [22] | Latent variable model with Lasso regularization | Produces positive-definite covariance matrices; enables sparse and interpretable inference | Limited to linear pairwise interactions; results sensitive to pseudo-count choice | counts = FALSE, pseudo = 0.5, k_cv = 3, lam_int = [1e-4, 1], k_max = 20, n_boot = 20 |
| CMIMN [15] | Conditional mutual information | Captures non-linear and conditional associations; suitable for complex microbial interactions | Assumes normality of data; only first-order CMI tests are considered due to computational intensity | quantitative = TRUE, q1 = 0.7, q2 = 0.95 |

### 2.5 Robustness evaluation via bootstrap analysis

To assess the reproducibility of `CMiNet` under sample-level variability and to determine an appropriate threshold for network construction, we performed a bootstrap-based evaluation. This analysis also allowed us to compare consensus networks with those generated by individual methods, which is particularly important in the absence of a gold standard for microbiome network inference. We generated 50 bootstrap replicates for each real dataset by resampling the data (i.e., rows) with replacement. Each replicate maintained the same number of samples as the original dataset, preserving the overall taxonomic distribution while introducing stochastic variation. This approach simulates the effect of analyzing alternative cohorts that share the same characteristics but differ due to natural sampling variability. For evaluation, we used the network inferred from the original dataset as the reference network for each method. Specifically, to evaluate the performance of a method *A*, we first inferred a network from the original dataset using method *A*; this network served as the gold standard to assess the accuracy of networks reconstructed from the bootstrap replicates using the*A* method. Each bootstrap network was then compared to its corresponding reference network using the F1-score, a metric that balances precision and recall to measure overall performance. We evaluated four inference strategies: 1- `CMiNet_All`: Consensus networks constructed using all available inference algorithms, with thresholds ranging from 0 to 9 (reflecting the use of up to all 10 methods); 2- `CMiNet`_Conditional: Consensus networks constructed using only conditional dependence-based methods, with thresholds ranging from 0 to 5 (up to 6 conditional dependence-based methods); 3- `CMiNet`_Correlation: Consensus networks constructed using only correlation-based methods, with thresholds ranging from 0 to 3 (up to 4 correlation-based methods); 4- Single Method: Networks constructed independently using each inference method without consensus. In addition, we computed 95% confidence intervals (CIs) for the F1-scores using bootstrap resampling [23]. The CIs were estimated using the percentile method, which involves calculating the 2.5th and 97.5th percentiles of the distribution of F1-scores across the 50 bootstrap replicates. Narrower confidence intervals indicate greater stability and suggest that the resulting networks are more reproducible and less sensitive to input variability. This analysis allowed us to evaluate the robustness of `CMiNet` and its reliability in reconstructing microbial interaction networks.

### 2.6 Identification of important taxa using multiple strategies

We emphasize the importance of constructing reliable microbiome networks not only to explore microbial interactions, but also as a foundation for identifying robust taxa associated with disease. One practical application of such networks—highlighted in this study—is identifying microbial biomarkers by comparing network roles across conditions. To identify taxa associated with scabpit disease, we applied three complementary strategies: *1. Machine Learning-Based Selection:* We normalized the microbiome data using total sum scaling (TSS) and applied several feature selection methods implemented in `scikit-learn` [24], including SelectKBest (based on ANOVA F-value), mutual information ranking, and Recursive Feature Elimination (RFE) using logistic regression, decision trees, gradient boosting, and random forests. Each taxon was assigned a score based on the number of different seven methods that selected it as important. We then computed the 85th percentile of these scores and retained taxa with scores equal to or greater than this threshold. These taxa were labeled as the *ML* set. *2. Differential Abundance Analysis:* We applied three widely used statistical tools to identify differentially abundant taxa: (1) Wilcoxon rank-sum test [25], (2) ALDEx2, which applies Welch’s t-test using Monte Carlo sampling, and [26](3) DESeq2, which models count data using the negative binomial distribution with FDR correction [27]. For each method, we sorted taxa by their FDR-adjusted *p*-values and selected those in the lowest 15%—corresponding to the most statistically significant results. This consistent thresholding approach allowed us to extract the top-performing taxa from each method, regardless of the specific statistical model it uses. The sets of important taxa resulting were labeled as *Wilcoxon, ALDEx2*, and *DESeq2*, respectively. 3. Network-Based Inference: We constructed microbial networks using `CMiNet` separately for healthy and diseased samples. To identify condition-specific important taxa, we computed Influence-Vertex Importance (IVI) scores [28] and compared their distributions across the two networks. Using multiple IVI-based strategies—including score thresholds, score differences, and rank shifts—we selected taxa with high centrality or major changes in influence, indicating potential biological relevance.

## 3 IMPLEMENTATION

The `CMiNet` Shiny app simplifies the process of constructing, visualizing, and analyzing microbiome networks through an interactive web-based platform. Unlike traditional R packages that require complex dependency installations, `CMiNet` provides an accessible and user-friendly interface, allowing researchers to upload data, select inference methods, visualize networks, and export results without prior programming expertise.

### 3.1 App Features and Functionality

The `CMiNet` Shiny app includes multiple tabs that guide users through the network construction and analysis process. In the *CMiNet Tab* users construct a weighted network by selecting inference algorithms and configuring algorithm-specific parameters. They can choose to apply all available algorithms or a subset based on their research needs. When the *Run* button is pressed, results for each selected algorithm are computed and made available in the *Results* sub-tab, where users can download individual algorithm outputs separately. Additionally, a consensus-weighted network is also generated and available for download. The *Visualization Tab* provides a comparative view of network structures based on four different thresholds. Each network displays the number of nodes with degree > 0, the number of edges, and the maximum degree value in the network. In the *Final Network Tab*, users can explore the final network structure by applying custom-defined thresholds. The *About Tab* provides detailed information on how to run `CMiNet` and interpret the results effectively.

### 3.2 Running CMiNet on the American Gut dataset

To demonstrate `CMiNet`’s functionality, we utilized the amgut1 dataset from the `SpiecEasi` R package [13], which is part of the American Gut Project—a large-scale citizen science initiative focused on the human gut microbiome. This dataset consists of 289 stool samples and 127 OTUs, and represents a well-preprocessed subset of human-associated microbial diversity. Figure 2 illustrates the interactive workflow of the `CMiNet` Shiny app and its application to the `amgut1.filt` dataset. After uploading the gut dataset, users can view a preview of the data in the *Data Head* sub-tab. Next, they select one or more inference algorithms and adjust algorithm-specific parameters. When the *Run* button is pressed, the app computes results for each selected method and displays them in the *Results* sub-tab to download. The *Visualization Tab* provides a comparative overview of networks constructed using four edge-weight thresholds. For instance, applying a threshold of 9 results in a sparse network with 55 nodes and 47 edges, while a threshold of 8 includes more edges and expands the network to 77 nodes and 94 edges. In the final step, users select a threshold (e.g., 9) to define their final microbiome network. Figure 3 illustrates the distribution of F1-scores across all methods based on 50 bootstrap replicates. Notably, `CMiNet` consistently outperformed individual methods, with thresholds 6, 7, and 8 yielding the highest average F1-scores and the lowest variability. This suggests that the consensus approach not only improves accuracy but also enhances the stability of microbial network inference. Combining all correlation-based methods also improved performance over individual correlation methods, while consensus networks constructed from five out of six conditional dependence-based methods significantly outperformed most single-method results. Among individual methods, some performed well on this dataset—such as `SpiecEasi_glasso` and `CCLasso`—but their performance may vary across datasets. In contrast, certain methods like SPRING exhibited lower F1-scores in this context, though they may perform better on other datasets. Additionally, methods like `Bicor` tended to produce dense networks with inflated true positive counts, but also a higher rate of false positives. At the other extreme, overly sparse networks generated at high thresholds (e.g., threshold 9) suffered from reduced recall, which led to lower overall F1-scores. Table S1 provides a detailed comparison of performance metrics across `CMiNet` consensus networks and individual network inference methods applied to the American Gut dataset. It includes the number of edges in the reference network, average number of inferred edges (± standard deviation) across 50 bootstrap replicates, mean true positives (TP), false positives (FP), precision, recall, and F1-score. This table reveals that increasing the `CMiNet` consensus threshold leads to sparser networks with fewer false positives, thereby enhancing network precision. As shown, intermediate thresholds (5–8) deliver the most balanced trade-off between precision and recall (up to 0.83), resulting in the highest F1-scores. In contrast, single-method networks often produce denser networks with more false positives and variable performance. These results demonstrate that `CMiNet`’s consensus-based filtering significantly improves both the robustness and interpretability of microbial network reconstruction. Figure S1 shows the 95% bootstrap confidence intervals for the F1-scores. As illustrated, `CMiNet` at thresholds 6 and 7 not only achieves high F1-scores but also exhibits narrow confidence intervals, indicating greater reliability and reproducibility across bootstrap replicates. These results underscore the suitability of `CMiNet` for robust microbiome network inference.

**Figure 2.**
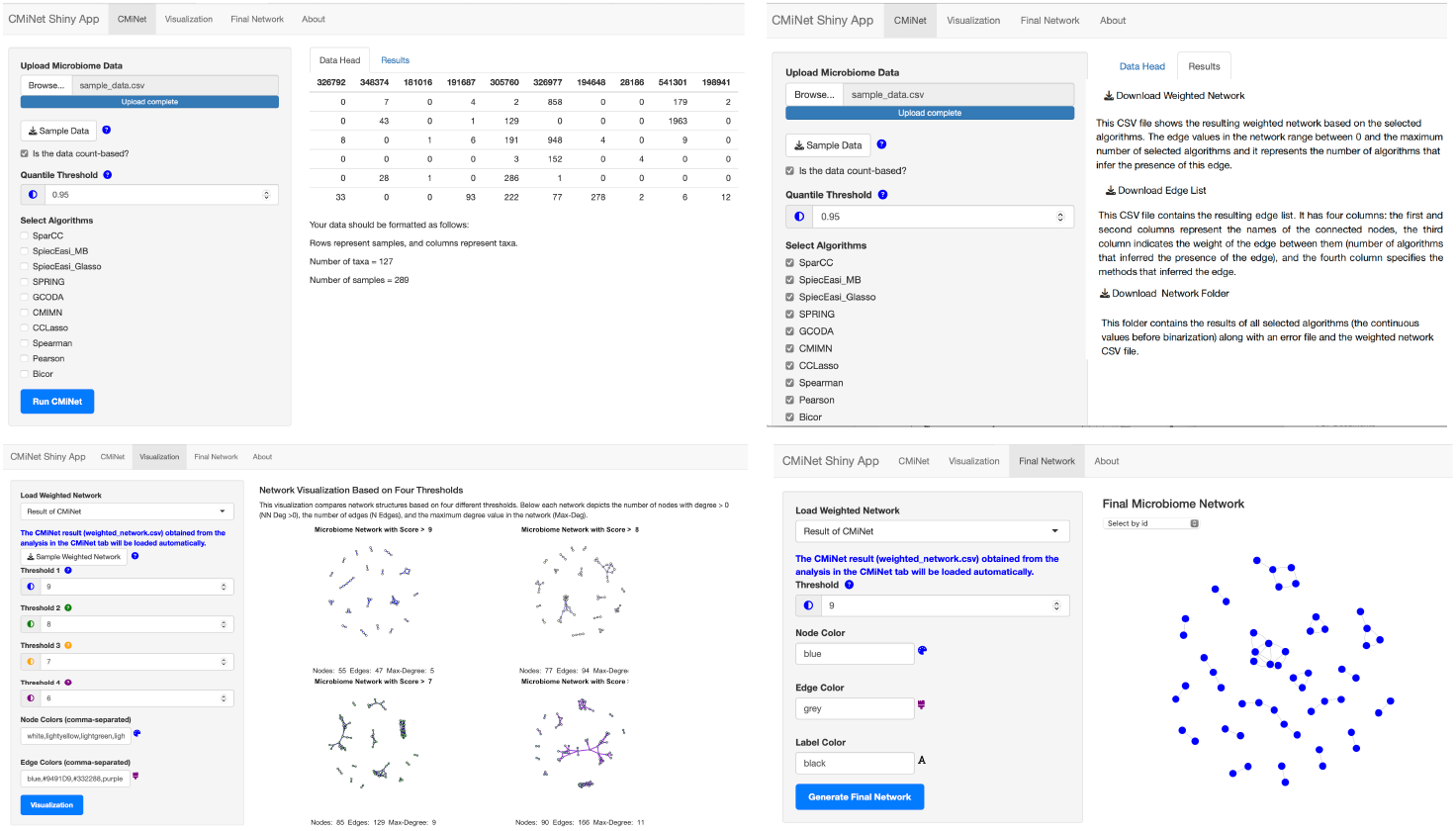
Overview of the CMiNet Shiny app interface and results based on the amgut1.filt dataset. (A) In the *CMiNet* tab, users upload microbiome abundance data, preview a summary in the *Data Head* sub-tab, and configure inference algorithms. Upon clicking *Run*, individual algorithm outputs and a consensus-weighted network are generated and displayed in the *Results* sub-tab. (B) The *Visualization* tab displays network structure across four edge-weight thresholds. For example, a threshold of 9 includes only edges supported by 9 or more algorithms, resulting in a sparser, high-confidence network (55 nodes, 47 edges). Lower thresholds yield denser networks. (C) In the *Final Network* tab, users can explore the final selected network (e.g., threshold 9), focusing on individual nodes and their neighbors.

**Figure 3.**
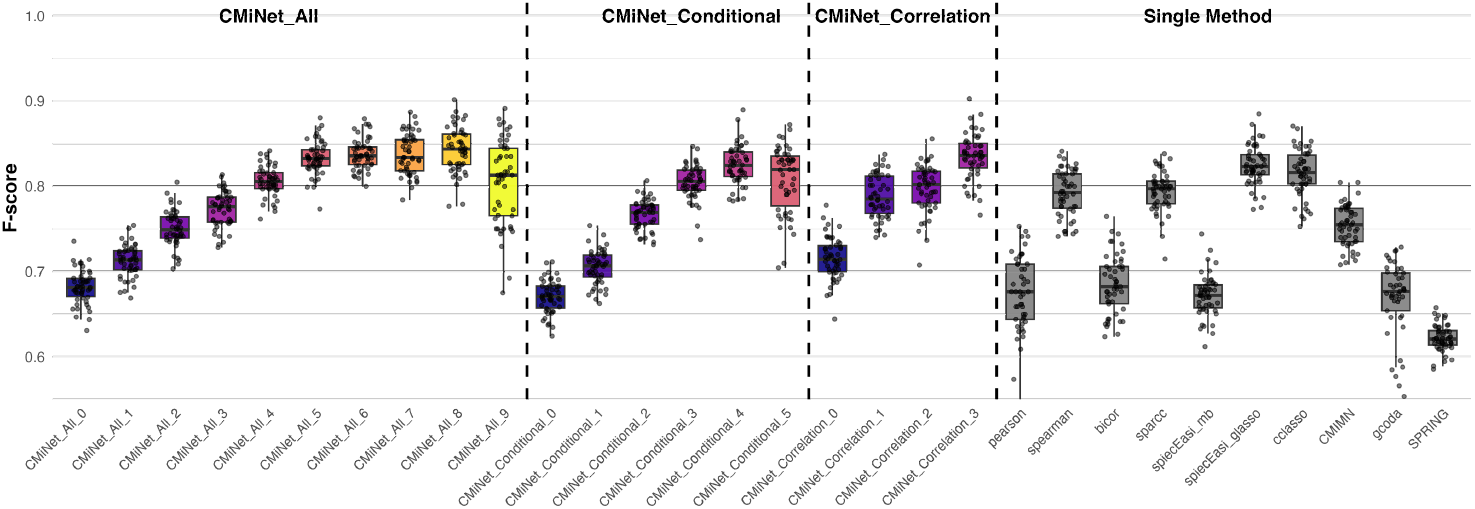
Robustness of microbial network inference across methods and CMiNet thresholds. F1-scores were computed for networks reconstructed from 50 bootstrap replicates using different strategies: CMiNet with thresholds from 0 to 9, correlation-based and conditional dependence-based consensus networks, and individual methods. Each dot represents the F1-score for a single bootstrap replicate, and boxplots summarize the distribution for each method. CMiNet_All at thresholds 6, 7, and 8 achieved the highest average F1-scores with lower variability, indicating improved robustness and accuracy compared to single-method approaches. Dashed vertical lines separate CMiNet_All, conditional-only, correlation-only, and single-method groups.

### 3.3 Running CMiNet on soil Microbiome data

We analyzed the soil microbiome at the Class taxonomic level using microbial abundance data derived from 256 pre-planting soil samples collected from potato fields in Minnesota (*n* = 108) and Wisconsin (*n* = 148). DNA was extracted and sequenced following the methods described in [8], capturing both bacterial and fungal communities associated with common scab disease. The 16S and ITS amplicon sequencing data are publicly available in the NCBI Short Read Archive under BioProject accession PRJNA1135141. At harvest, disease severity was recorded as a binary outcome where tubers with pitted scab lesions were labeled as diseased (1) and those without lesions as healthy (0). We filtered the dataset to retain only taxa present in at least 15 samples, and samples with at least 20 non-zero taxa and a total abundance of 20 or more. After filtering, 214 samples remained (115 healthy and 99 diseased), with a total of 99 microbial classes. To identify an appropriate threshold for constructing soil microbiome networks—similar to the approach used for the `amgut1.filt` dataset—we performed a bootstrap-based robustness analysis. The results are shown in Figure 4 and Figure S2. As shown in Table S2, the performance of `CMiNet` on the soil microbiome dataset demonstrates the benefits of consensus network construction. At intermediate thresholds (particularly 5–7), `CMiNet` achieves high F1-scores (up to 0.8), reflecting a favorable balance between precision and recall. These thresholds retain enough edges to capture true microbial interactions while effectively filtering out false positives. In contrast, single-method approaches, such as `SPRING` and `gcoda`, exhibit higher variability and lower precision, often leading to inflated networks. Notably, combining correlation-based methods alone also yields competitive results, but the best performance is observed when integrating multiple inference strategies through consensus filtering. These results reinforce the importance of robust network construction in identifying biologically meaningful microbial interactions. To identify important taxa from networks, we constructed microbial networks using `CMiNet` separately for healthy and diseased samples. We applied a consensus threshold of 7 (i.e., retaining edges supported by at least 8 out of 10 methods). This threshold produced networks that were neither overly sparse nor excessively dense, while also achieving high F1-scores and narrow confidence intervals—indicating both accuracy and reproducibility. The resulting healthy network was denser, containing 58 edges, compared to only 44 in the diseased network, suggesting a disruption in microbial connectivity under disease conditions. In each network, we computed IVI scores, and the distribution of IVI scores showed a stark contrast between conditions. The 85th percentile of IVI scores was 16.53 for the healthy network and only 1.19 for the diseased network, further highlighting the loss of connectivity in disease network. Based on IVI scores, we defined four sets of important taxa to capture different aspects of taxon centrality and condition-specific influence. *IVI_healthy*: This set includes taxa whose IVI scores in the healthy network are above the 85th percentile (16.5263). *IVI_disease*: This set consists of taxa with IVI scores in the disease network above the 85th percentile (1.192884) that are not also included in IVI_healthy. *IVI_difference*: Taxa in this set exhibit the largest absolute differences in IVI scores between the healthy and disease networks, indicating major shifts in influence. *IVI_rank_difference*: This set captures taxa with the most significant changes in IVI ranking positions between networks, highlighting shifts in relative importance that might not be reflected in score magnitude alone. Figure 5 displays intersections among different eight strategies to select disease related taxa using an UpSet plot. Some taxa were identified through machine learning or differential abundance methods, while others were derived from network-based inference. To evaluate agreement among the eight strategies (*ML, Wilcoxon, ALDEx2, DESeq2, IVI_healthy, IVI_disease, IVI_difference, IVI_rank_difference*), we computed Jaccard-based percentage overlap (Figure S3). For instance, ML and DESeq2 shared 45.8% overlap, while ML and IVI_difference shared 20.7%. To assess the consistency of important taxa identified by different strategies, we constructed a binary presence matrix indicating which taxa were selected as important by each of the eight methods: ML-based feature selection, Wilcoxon, ALDEx2, DESeq2, IVI_healthy, IVI_disease, IVI_difference, and IVI_rank_difference. The heatmap (Figure S4) visualizes both method-specific selections and areas of overlap, with highly consistent taxa appearing in the upper rows. This comprehensive comparison emphasizes the value of integrating multiple analytical strategies to identify robust disease-associated taxa, and underscores the benefit of network-based features in complementing traditional differential abundance approaches.

**Figure 4.**
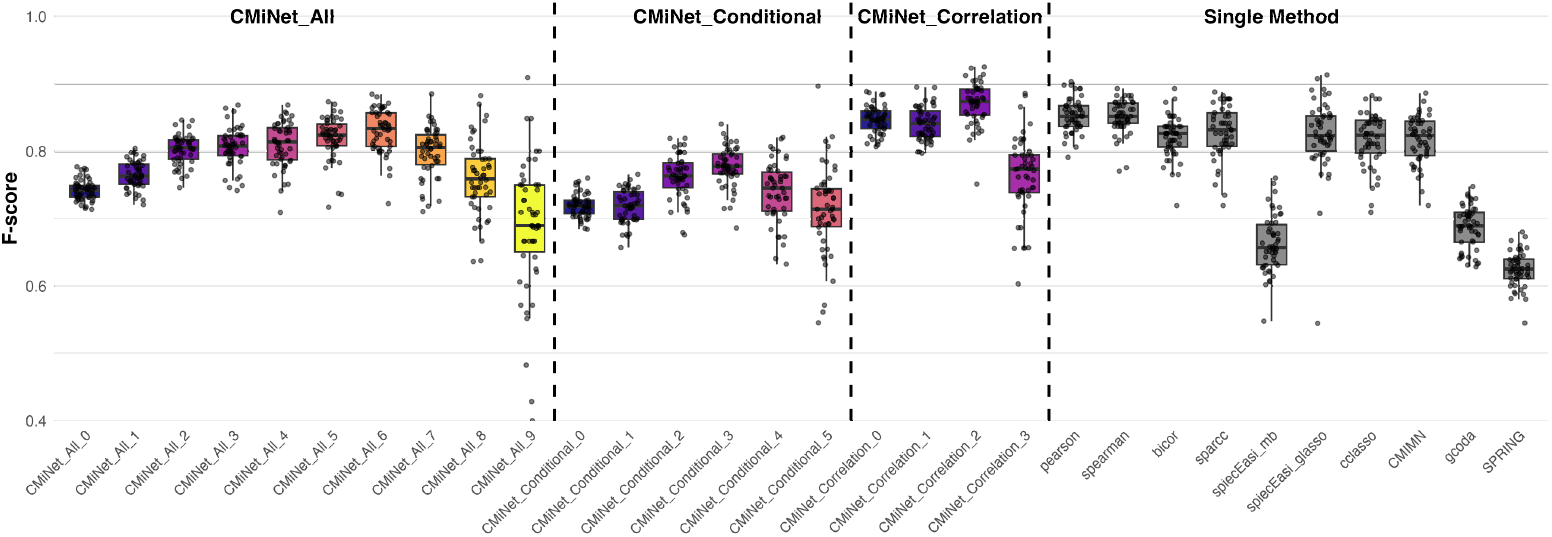
Robustness of microbial network inference across methods and CMiNet thresholds. F1-scores were computed for networks reconstructed from 50 bootstrap replicates using different strategies: CMiNet with thresholds from 0 to 9, correlation-based and conditional dependence-based networks, and individual methods. Each dot represents the F1-score for one bootstrap replicate, and boxplots summarize the distribution per method. CMiNet at thresholds 6 and 7 achieved the highest average F1-scores with lower variability, demonstrating improved robustness and accuracy compared to single methods. Dashed vertical lines separate CMiNet thresholds, conditional methods, correlation-based methods, and single-method baselines.

**Figure 5.**
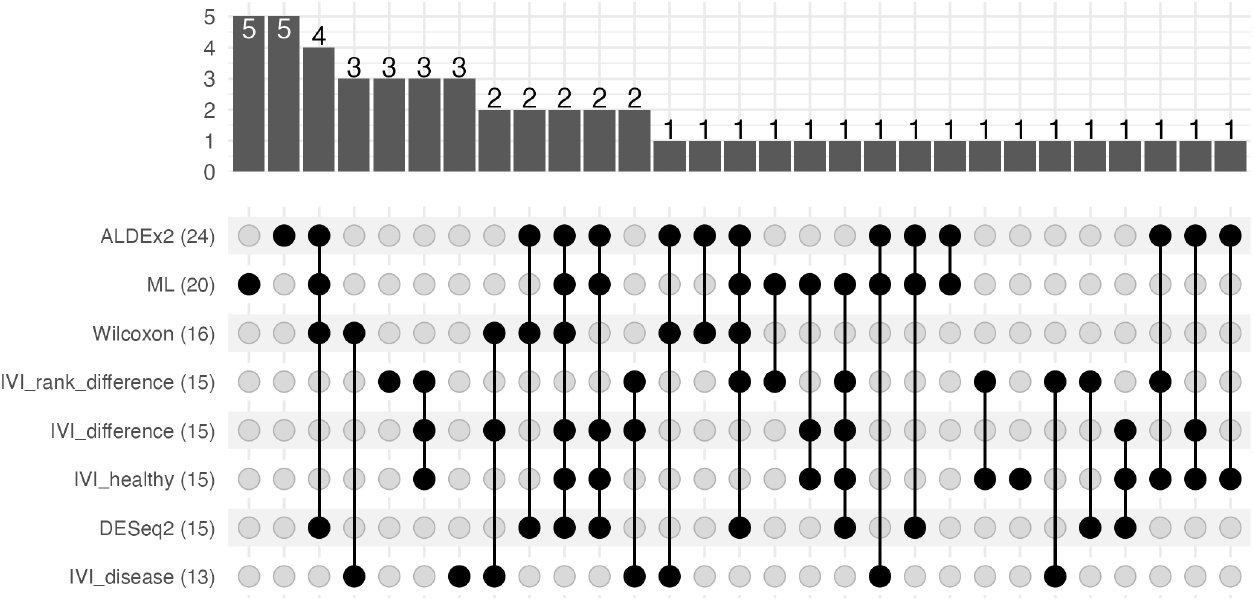
UpSet plot showing the intersections among eight strategies used to identify important microbial taxa. The methods (rows) include machine learning (ML), three differential abundance techniques (ALDEx2, Wilcoxon, DESeq2), and four network-based approaches (IVI_difference, IVI_rank_difference, IVI_healthy, IVI_disease). The method names are displayed on the left, each followed by the number of taxa it identified (set size). The vertical bars represent the number of taxa shared by the corresponding combination(s) of methods marked below each bar. For example, 5 taxa were uniquely identified by ML, 5 only by ALDEx2, and 4 taxa were jointly identified by ML, ALDEx2, Wilcoxon, and DESeq2 strategies.

Importantly, *Ktedonobacteria, Acidobacteriae, Vicinamibacteria, MB-A2-108, Planctomycetes*, and *Anaerolineae* were consistently identified as important across multiple independent analyses. Specifically, they were selected by five or six out of the eight distinct strategies applied, including machine learning-based feature selection, three differential abundance tests (Wilcoxon, ALDEx2, DESeq2), and four network-based inference approaches based on IVI scores. This strong cross-method support highlights their robust association with the disease condition in our study and underscores their potential biological significance. *Ktedonobacteria* are an ancient, spore-forming bacterial class widely distributed across terrestrial environments, including soil, bark, and sand, and they persist even under extreme conditions such as geothermal areas. Their large genome size, complex life cycle, and high biosynthetic potential for novel secondary metabolites make them a promising microbial resource for natural product discovery and antimicrobial activity [29, 30]. *Acidobacteriae* and *Vicinamibacteria* were identified as enriched bacterial classes in the BC site based on LEfSe analysis of rhizosphere soils. Their enrichment suggests a potential role in shaping microbial community composition under specific environmental or agricultural conditions [31]. The bacterial class *MB-A2-108*, part of the phylum Actinobacteria, exhibited high relative abundance in soils conducive to Prunus replant disease (PRD), suggesting its potential involvement in soilborne disease processes. Furthermore, *MB-A2-108* significantly increased following afforestation, indicating that it is a responsive and ecologically important taxon potentially contributing to soil health, nutrient cycling, and structural changes under both natural and disturbed environmental conditions [32, 33]. *Planctomycetes* are a widespread and diverse bacterial phylum abundant in soils across five continents, with community composition strongly shaped by soil management history, organic matter content, and pH. They also play a critical role in ecosystem processes, including the decomposition of Sphagnum-derived litter, underscoring their importance in nutrient cycling and organic matter turnover [34, 35]. *Anaerolineae* are strictly anaerobic, fermentative bacteria capable of utilizing a broad range of substrates, and have been associated with waterlogged agricultural soils and nitrogen cycling processes. Their increased abundance in response to pathogen burden, irrigation, tillage, and nitrogen fertilization highlights their role in shaping soil health and regulating biogeochemical nutrient dynamics [29, 36].

## 4 Conclusion

Understanding microbial interactions is essential for revealing how communities function, maintain stability, and respond to environmental changes. Beyond these ecological insights, we designed a comparative analysis of microbial networks in diseased and healthy conditions to investigate how interaction patterns shift in response to disease and to identify condition-specific taxa with potential as biomarkers. Although many algorithms have been developed to infer microbiome networks, significant variation among inference methods and the absence of a gold standard for validation make network-based microbiome analysis challenging and often irreproducible. `CMiNet` addresses these challenges by offering a flexible, user-friendly platform that integrates ten established network inference algorithms. It enables users to construct microbial networks based on correlation-based methods, conditional independence-based methods, or a combination of both. This structured framework allows researchers to tailor network construction based on their specific goals—whether prioritizing co-occurrence patterns, direct interactions, or consensus across methods. A key strength of `CMiNet` is its consensus-based filtering, which retains only interactions supported by multiple algorithms, thereby minimizing method-specific biases and enhancing reproducibility. This approach leads to more robust microbial networks that better reflect underlying biology. The app’s interactive interface simplifies the entire process—from data upload and algorithm selection to visualization and export—making advanced microbial network analysis accessible to researchers without programming expertise. We demonstrated the effectiveness of `CMiNet` on both human gut and soil microbiome datasets. In the gut microbiome case, we used bootstrap analysis to compare the robustness of `CMiNet` against individual methods, showing that consensus networks achieved higher F1-scores and narrower confidence intervals. In the soil microbiome analysis, we began by determining a reliable threshold for consensus network construction through a robustness evaluation across healthy and diseased samples. The resulting networks differed in their structure— the healthy network was more densely connected, while the diseased network showed reduced connectivity, indicating potential disruption in microbial interactions. To identify disease-associated taxa, we designed a multistep approach that combined diverse strategies, including machine learning, differential abundance analysis, and network-based scoring. This comparative analysis of healthy and diseased networks revealed condition-specific central taxa, demonstrating how robust network construction enhances the accuracy of biomarker discovery. Importantly, `CMiNet` does not merge fundamentally different network types blindly. Instead, it empowers users to explore each method class independently before generating a consensus network that retains only highly supported edges, ensuring biological interpretability and statistical rigor. Users also have access to detailed information about which specific methods support each edge, enabling them to filter results based on trusted or context-relevant algorithms. This level of transparency and customization makes `CMiNet` a powerful and adaptable tool for a wide range of microbiome research applications and study designs. Building on these advances, our integrative strategy also identified a set of robust taxa consistently associated with the disease condition across multiple independent methods. Specifically, *Ktedonobacteria, Acidobacteriae, Vicinamibacteria, MB-A2-108, Planctomycetes*, and *Anaerolineae* were selected by five or six out of eight distinct feature selection strategies, underscoring their potential biological relevance. These findings highlight how combining consensus network construction with complementary statistical approaches can not only improve network reliability but also facilitate the discovery of key microbial indicators linked to health and disease states.

## Data availability statement

The `amgut1.filt` dataset used in this study is publicly available and included in the `SpiecEasi` R package. For convenience, it is also bundled with the `CMiNet` R package as a demonstration dataset to facilitate reproducibility and user testing. The soil microbiome dataset analyzed in this study is available through the NCBI Short Read Archive under BioProject accession number PRJNA1135141, as described in [8]. All source code, including the R package and Shiny app, along with documentation and reproducible examples, is freely available on GitHub: R package: https://github.com/solislemuslab/CMiNet Shiny App: https://github.com/solislemuslab/CMiNetShinyAPP Public version of the app: https://cminet.wid.wisc.edu.

## Acknowledgements

We thank Dr. Richard Lankau and Dr. Shan Shan for providing access to the soil microbiome data used in this study, available through the NCBI Short Read Archive under BioProject accession number PRJNA1135141. We also thank Benjamin Huebner at the University of Wisconsin–Madison Research Computing Services for assisting with the deployment of the public version of the `CMiNet` Shiny app. Finally, we thank Dr. Maryam Shahdoost for checking, installing, and running the `CMiNet` package.

## SUPPLEMENTARY MATERIAL: AN R PACKAGE AND USER-FRIENDLY SHINY APP FOR CONSTRUCTING CONSENSUS MICROBIOME NETWORKS

**Figure 1.**
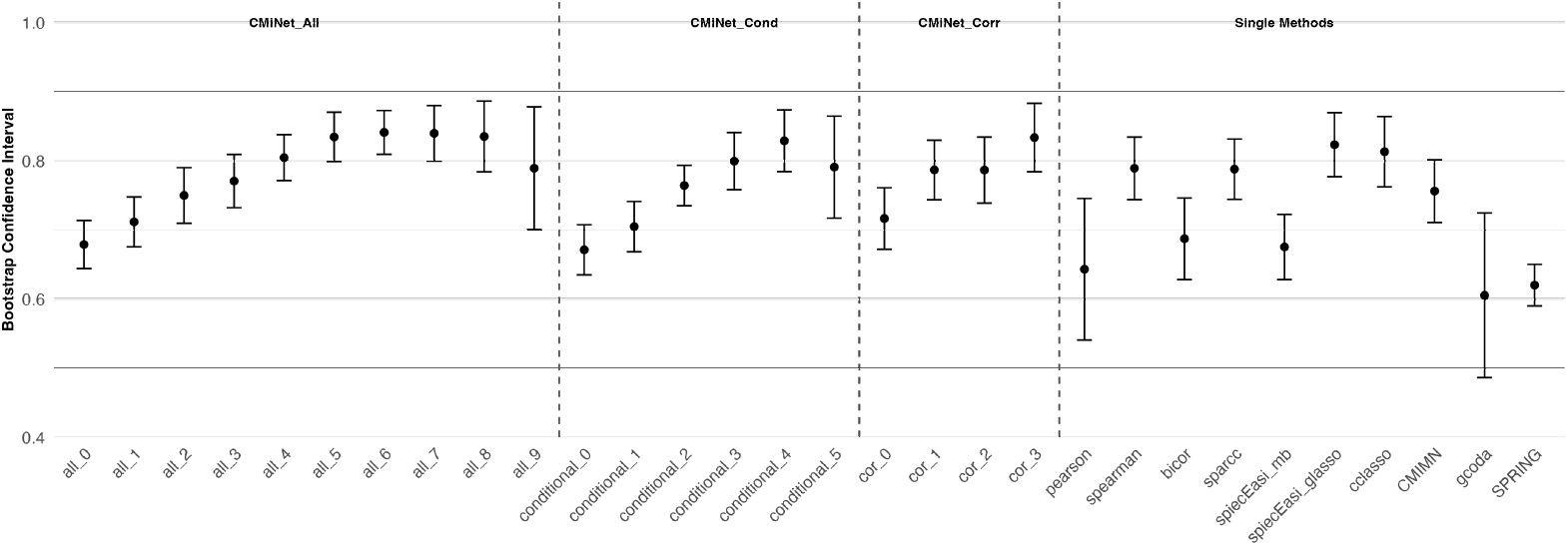
Bootstrap confidence intervals of F1-scores for each network inference method. The plot displays the 95% bootstrap confidence intervals for F1-scores across all methods, providing a measure of variability and reliability in microbial network reconstruction. CMiNet_All with threshold 6 exhibits the narrowest confidence intervals and consistently high F1-scores, indicating stable and reproducible performance across bootstrap replicates. In contrast, single-method approaches show wider confidence intervals, reflecting greater sensitivity to input variability and reduced robustness.

**Figure 2.**
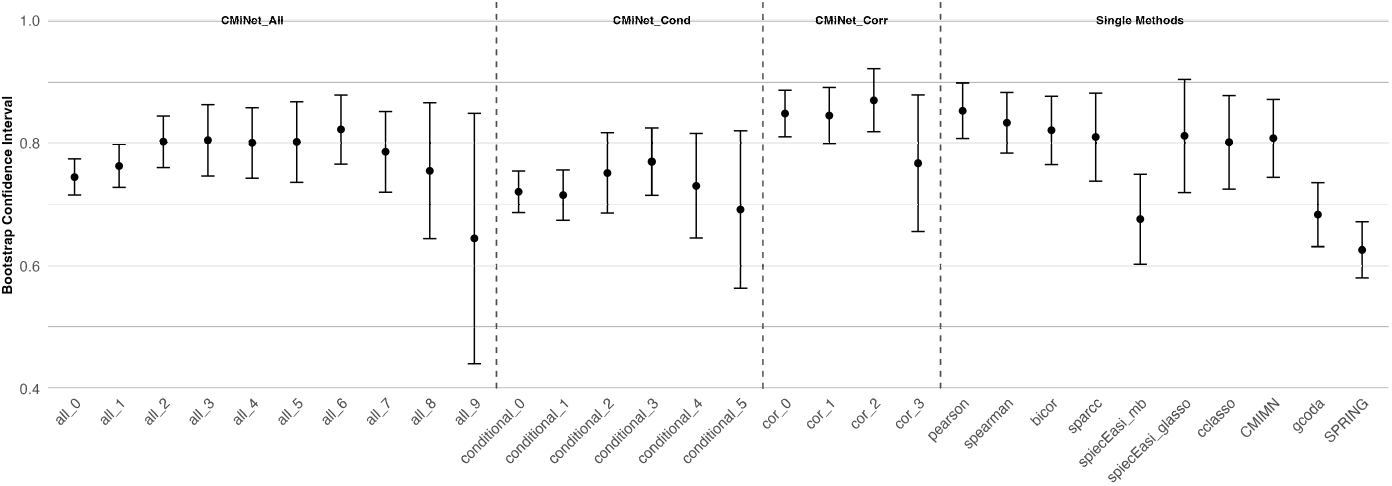
Bootstrap confidence intervals of F1-scores for each network inference method. The plot displays the 95% bootstrap confidence intervals for F1-scores across all methods, providing a measure of variability and reliability in microbial network reconstruction. CMiNet thresholds 6 and 7 show narrow confidence intervals with consistently high F1-scores, indicating stable and reproducible performance across bootstrap replicates. In contrast, single-method approaches exhibit wider confidence intervals, reflecting greater sensitivity to input variability and reduced robustness.

**Figure 3.**
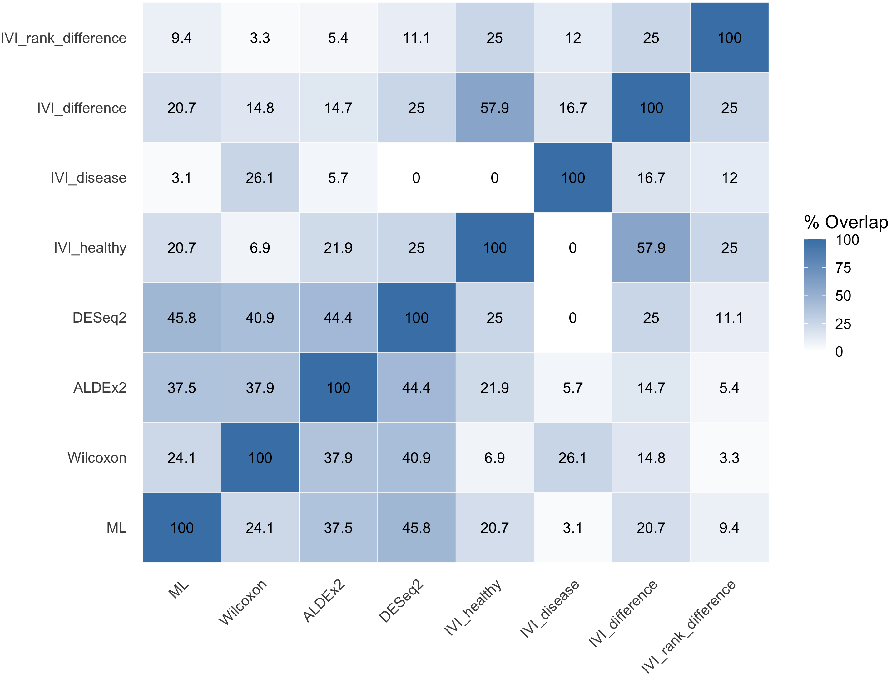
Heatmap showing Jaccard-based percentage overlap between important taxa identified by eight different methods.

**Figure 4.**
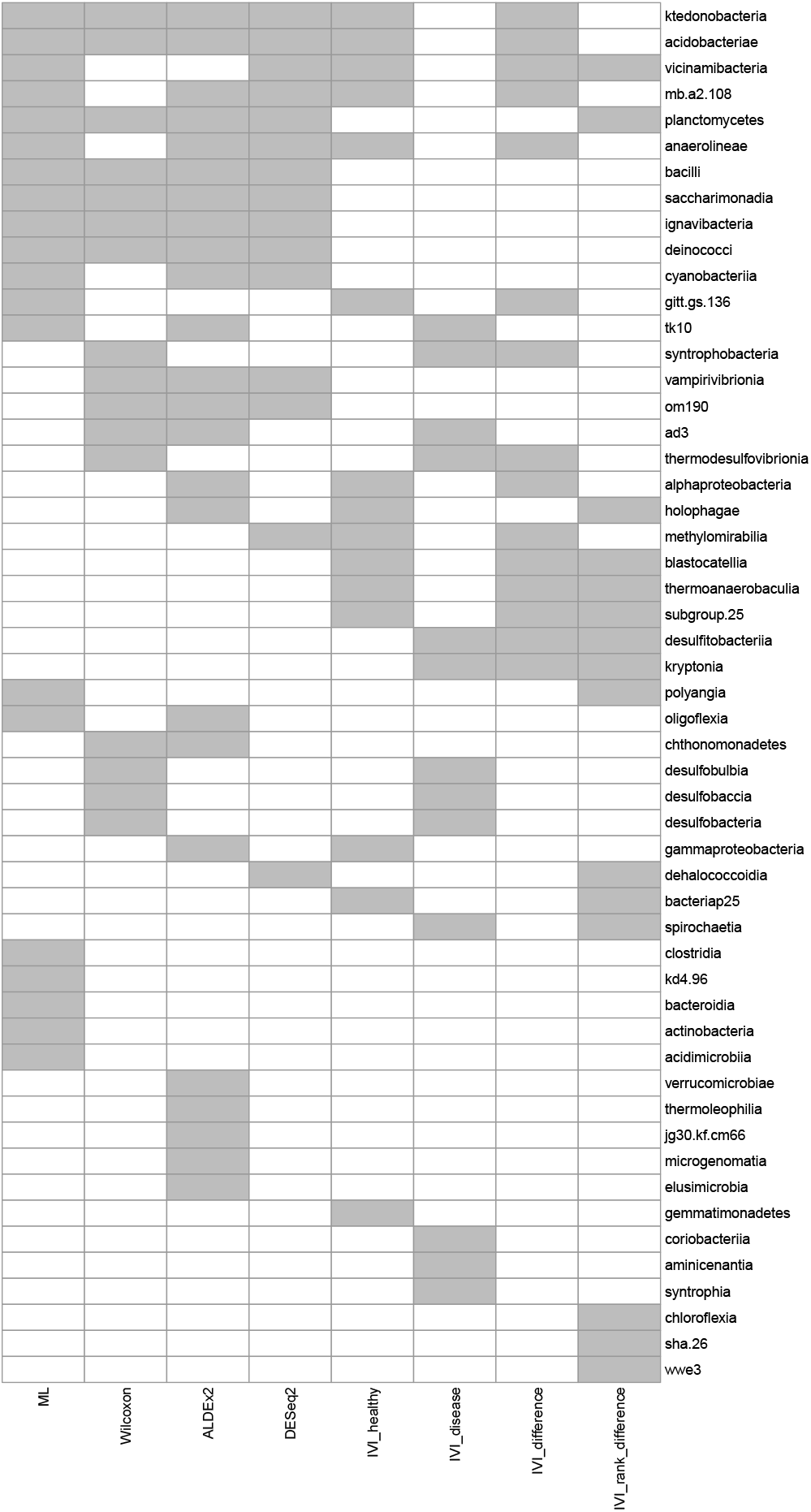
Heatmap of important taxa identified by eight selection strategies. Each row represents a taxon, and each column corresponds to a selection method: machine learning (ML), three differential abundance techniques (Wilcoxon, ALDEx2, DESeq2), and four network-based approaches (IVI_healthy, IVI_disease, IVI_difference, IVI_rank_difference). Gray cells indicate that the taxon was selected as important by the corresponding method. Taxa are sorted by the number of supporting methods, with those confirmed by multiple approaches at the top, highlighting robust and reproducible taxa selection across methodologies.

**Table 1.**
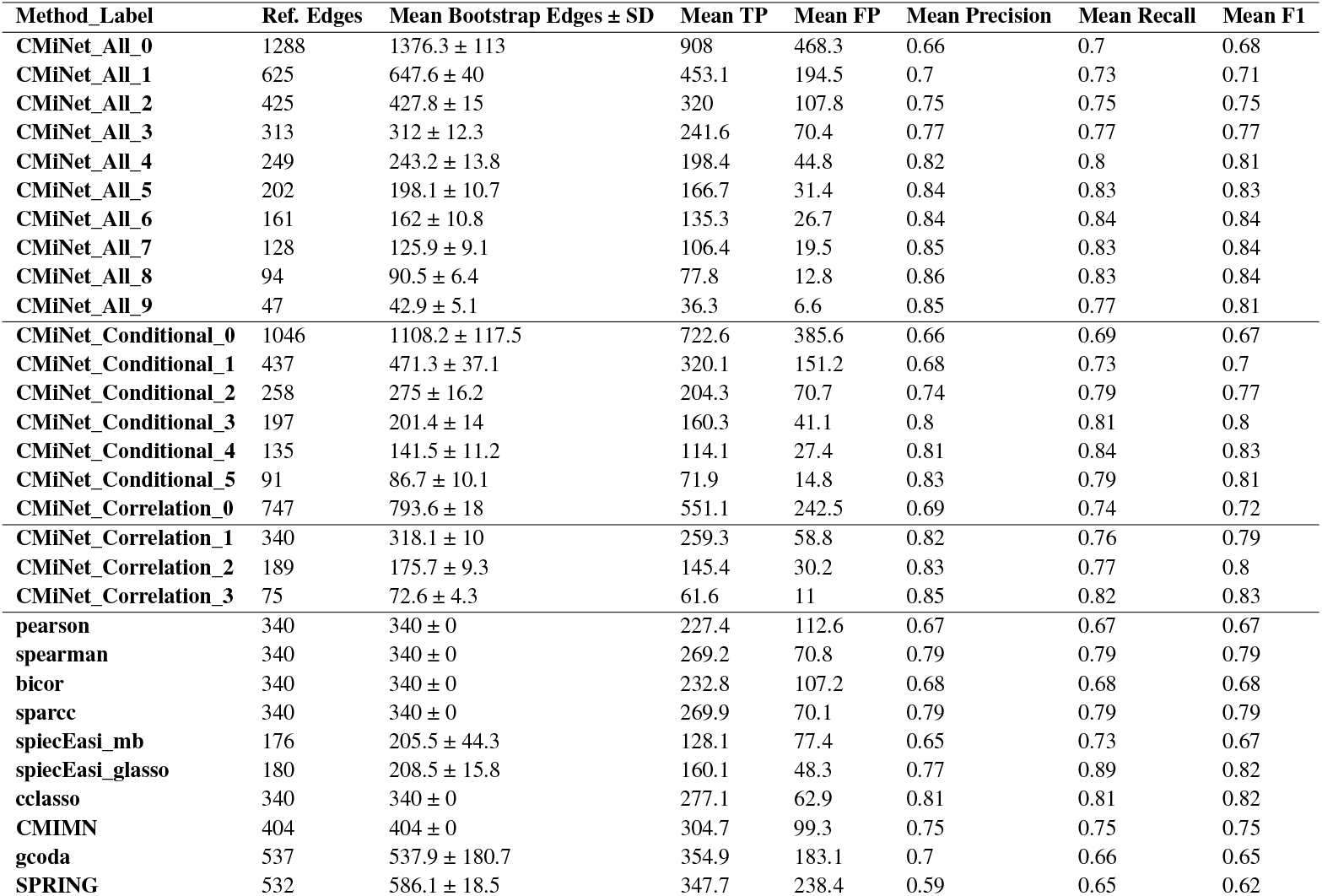
Performance summary of CMiNet and individual methods in microbial network inference on the American Gut dataset. For each method or consensus threshold, we report the number of edges in the reference network, mean number of edges inferred from 50 bootstrap replicates (± standard deviation), mean true positives (TP), false positives (FP), precision, recall, and F1-score. `CMiNet` consensus networks at intermediate thresholds (especially thresholds 6–8) achieved high precision and recall, indicating robust performance. In contrast, single-method networks exhibited more variability in precision and recall, reflecting method-specific biases.

**Table 2.**
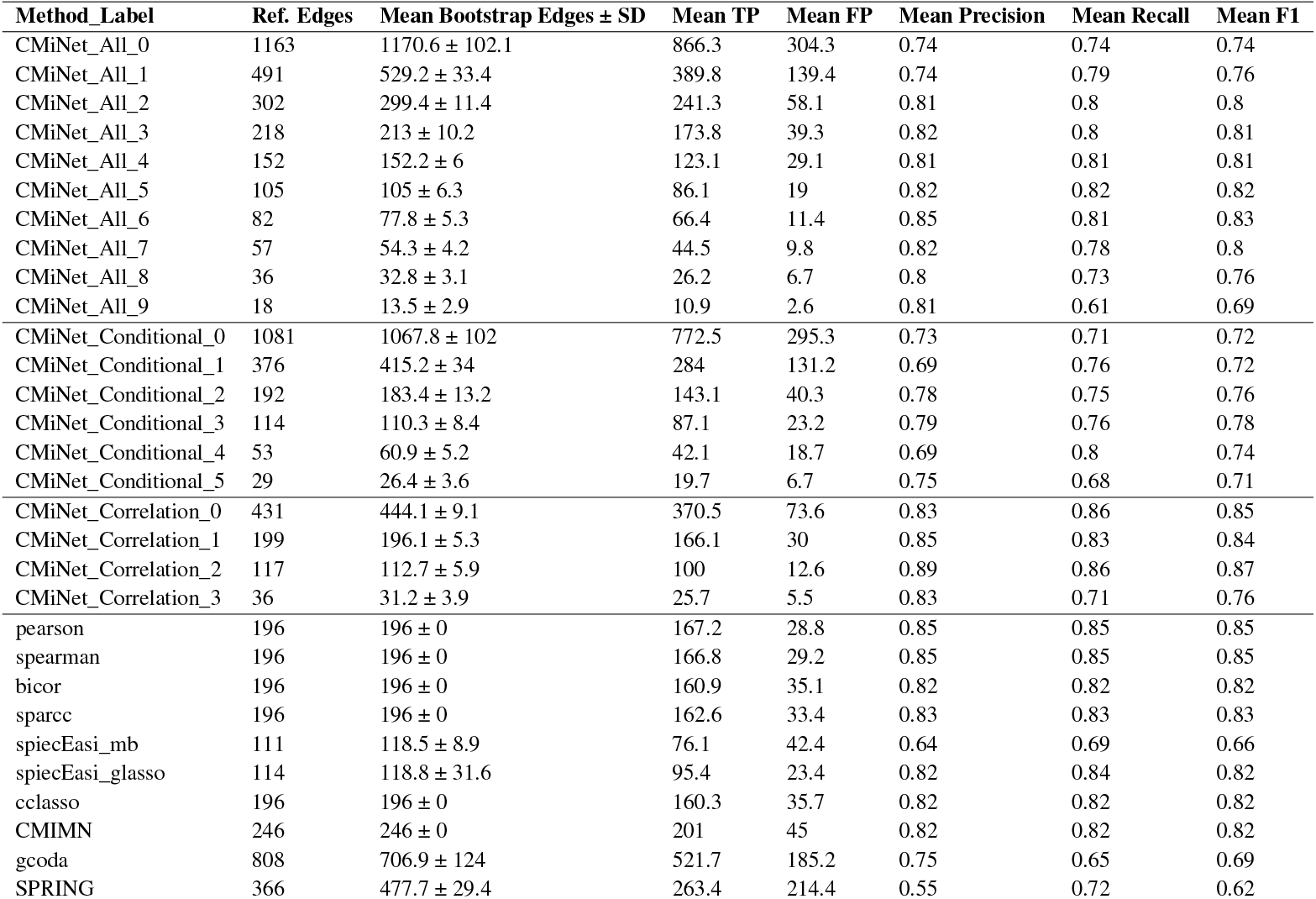
Performance summary of CMiNet and individual methods in microbial network inference applied to the soil microbiome dataset. This table presents the average performance across 50 bootstrap replicates for each method. Metrics include the number of edges in the reference network (Ref. Edges), mean number of inferred edges ± standard deviation (Mean Bootstrap Edges), mean true positives (TP), mean false positives (FP), and mean values for precision, recall, and F1-score. *CMiNet* consensus networks at intermediate thresholds (e.g., 5–7) achieve a strong balance between precision and recall, resulting in high F1-scores and improved robustness compared to both lower thresholds (which yield overly dense networks) and higher thresholds (which may under-detect true interactions). Single-method approaches show greater variability and often higher false positive rates.

| Method_Label | Ref. Edges | Mean Bootstrap Edges ± SD | Mean TP | Mean FP | Mean Precision | Mean Recall | Mean F1 |
| --- | --- | --- | --- | --- | --- | --- | --- |
| CMiNet_All_0 | 1163 | 1170.6 ± 102.1 | 866.3 | 304.3 | 0.74 | 0.74 | 0.74 |
| CMiNet_All_1 | 491 | 529.2 ± 33.4 | 389.8 | 139.4 | 0.74 | 0.79 | 0.76 |
| CMiNet_All_2 | 302 | 299.4 ± 11.4 | 241.3 | 58.1 | 0.81 | 0.8 | 0.8 |
| CMiNet_All_3 | 218 | 213 ± 10.2 | 173.8 | 39.3 | 0.82 | 0.8 | 0.81 |
| CMiNet_All_4 | 152 | 152.2 ± 6 | 123.1 | 29.1 | 0.81 | 0.81 | 0.81 |
| CMiNet_All_5 | 105 | 105 ± 6.3 | 86.1 | 19 | 0.82 | 0.82 | 0.82 |
| CMiNet_All_6 | 82 | 77.8 ± 5.3 | 66.4 | 11.4 | 0.85 | 0.81 | 0.83 |
| CMiNet_All_7 | 57 | 54.3 ± 4.2 | 44.5 | 9.8 | 0.82 | 0.78 | 0.8 |
| CMiNet_All_8 | 36 | 32.8 ± 3.1 | 26.2 | 6.7 | 0.8 | 0.73 | 0.76 |
| CMiNet_All_9 | 18 | 13.5 ± 2.9 | 10.9 | 2.6 | 0.81 | 0.61 | 0.69 |
| CMiNet_Conditional_0 | 1081 | 1067.8 ± 102 | 772.5 | 295.3 | 0.73 | 0.71 | 0.72 |
| CMiNet_Conditional_1 | 376 | 415.2 ± 34 | 284 | 131.2 | 0.69 | 0.76 | 0.72 |
| CMiNet_Conditional_2 | 192 | 183.4 ± 13.2 | 143.1 | 40.3 | 0.78 | 0.75 | 0.76 |
| CMiNet_Conditional_3 | 114 | 110.3 ± 8.4 | 87.1 | 23.2 | 0.79 | 0.76 | 0.78 |
| CMiNet_Conditional_4 | 53 | 60.9 ± 5.2 | 42.1 | 18.7 | 0.69 | 0.8 | 0.74 |
| CMiNet_Conditional_5 | 29 | 26.4 ± 3.6 | 19.7 | 6.7 | 0.75 | 0.68 | 0.71 |
| CMiNet_Correlation_0 | 431 | 444.1 ± 9.1 | 370.5 | 73.6 | 0.83 | 0.86 | 0.85 |
| CMiNet_Correlation_1 | 199 | 196.1 ± 5.3 | 166.1 | 30 | 0.85 | 0.83 | 0.84 |
| CMiNet_Correlation_2 | 117 | 112.7 ± 5.9 | 100 | 12.6 | 0.89 | 0.86 | 0.87 |
| CMiNet_Correlation_3 | 36 | 31.2 ± 3.9 | 25.7 | 5.5 | 0.83 | 0.71 | 0.76 |
| pearson | 196 | 196 ± 0 | 167.2 | 28.8 | 0.85 | 0.85 | 0.85 |
| spearman | 196 | 196 ± 0 | 166.8 | 29.2 | 0.85 | 0.85 | 0.85 |
| bicor | 196 | 196 ± 0 | 160.9 | 35.1 | 0.82 | 0.82 | 0.82 |
| sparcc | 196 | 196 ± 0 | 162.6 | 33.4 | 0.83 | 0.83 | 0.83 |
| spiecEasi_mb | 111 | 118.5 ± 8.9 | 76.1 | 42.4 | 0.64 | 0.69 | 0.66 |
| spiecEasi_glasso | 114 | 118.8 ± 31.6 | 95.4 | 23.4 | 0.82 | 0.84 | 0.82 |
| cclasso | 196 | 196 ± 0 | 160.3 | 35.7 | 0.82 | 0.82 | 0.82 |
| CMIMN | 246 | 246 ± 0 | 201 | 45 | 0.82 | 0.82 | 0.82 |
| gcoda | 808 | 706.9 ± 124 | 521.7 | 185.2 | 0.75 | 0.65 | 0.69 |
| SPRING | 366 | 477.7 ± 29.4 | 263.4 | 214.4 | 0.55 | 0.72 | 0.62 |

